# Integrated Network Pharmacology and Experimental Analysis Unveil Modulation of EGFR/MAPK Signaling Cascades in Acute Cerebral Ischemia-Reperfusion Injury by Qing-Tong-Hua-Yu Decoction

**DOI:** 10.1101/2022.11.04.515245

**Authors:** Jiajing Hu, Long Zuo, Wenyu Qu, Hongdun He, Jie Bao, Wenyan Zhang, Yunyang Zhang, Meizhen Zhu, Tian Li

## Abstract

**Objective:** Based on network pharmacology, the response of Qing-tong-hua-yu Decoction (QTHY) to the regulation of EGFR/MAPK signaling cascade in cerebral ischemia-reperfusion injury was discussed and the possible mechanism of the protective effect of QTHY on the cerebral ischemia-reperfusion injury was studied.

**Methods:** A compound-target disease-function-pathway network was established and analyzed based on the network pharmacology approach used in Chinese medicine. The correlation, which is between effect of the components of QTHY Decoction against CI/RI with EGFR/MAPK signalling cascade response, was observed. And then the degree of neurological deficits in each group was assessed after cerebral ischemia for 2 hours and reperfusion for 3 hours, 24 hours, 3 days and 7 days. Expression levels of EGFR and p44/42MAPK in ischemic brain tissue at different time points in various groups of rats were tested by Western bolt (WB), real-time quantitative PCR (RT-qPCR) and immunohistochemistry (IHC).

**Results:** Network pharmacology analysis revealed that QTHY-mediated treatment involved 439 key targets, in which the effect of QTHY groups against CI/RI was associated with EGFR/MAPK signaling cascade. QTHY treatment reduced neurological deficit scores and improved ischemic changes in rats. In addition, QTHY promoted EGFR and p44/42MAPK expression in the SVZ through the EGFR/MAPK signaling cascade, with varying degrees of improvement at different time points.

**Conclusion:** QTHY can better improve cerebral ischemia injury in CI / RI rats and exert the neuroprotective effect of cerebral ischemia-reperfusion injury. This may be related to the potential of QTHY to activate the EGFR / MAPK signaling cascade, which is consistent with the results of network pharmacology analysis.

## 1. Introduction

Cerebral ischemic stroke (CIS), due to disturbance of cerebral circulation in a short time which leads brain cells to be ischemic, hypoxic or necrotic, is a kind of disease which can cause hemiplegia, speech problems, and even coma in clinical situation due to neurological deficits.

In order to improve or restore blood perfusion in the ischaemic area, the current clinical treatment of ischaemic stroke relies on activating blood vessels and inhibiting platelet aggregation primarily. However, this approach frequently results in cerebral ischemia-reperfusion injury (CI/RI), which is a significant contributor to the high disability and mortality rates of ischaemic stroke[1]. According to research, MAPK signaling pathway play a crucial role on the progression of cerebral ischaemia. After cerebral ischemia and hypoxia, MAPK signaling pathway which is found in central nervous system is activated and produces a important effect to control survival and apoptosis of neurons[2].At the same time, signaling pathway controlled by epidermal growth factor (EGF) play a pivotal role on promoting neurogenesis[3-5]. Besides, EGF is one of specific ligand of the epidermal growth factor receptor (EGFR). EGFR combining with EGF can activates the expression of receptor tyrosine kinases (RTKs), and then affects cell proliferation, differentiation, migration or metabolic changes. Furthermore, EGFR and fibroblast growth factor receptor (FGFR) synergistically regulate the ERK/MAPK pathway[6]. Promoting the expression of EGFR can help brain cells be restored and MAPK pathway be improved effectively. In a word, EGFR/MAPK signaling cascade is a crucial therapeutic target for the activation of neurogenesis and brain repair in the treatment of post stroke disability.

The fundamental pathogenesis of ischemic stroke is the overlap of pathogenic elements such stasis, phlegm, heat, and toxicity, which causes “paralysis of the cerebral veins.” Approaches to treat ischemic stroke should focus on “unblocking the cerebral veins and replenishing the cerebral marrow”, to unblocking the cerebral veins likes clear away blood stasis and heat, clear away phlegm and toxin, or clear up brain collaterals. Based on those approaches, our group had chosen Qing-tong-hua-yu Decoction (QTHY) to treat ischaemic stroke of the phlegm-stasis-heat knot type in clinical practice and discovered that it can successfully lessen clinical symptoms and hasten the recovery of neurological deficits. So we think it can protect brain tissue cells and lessen cerebral ischaemia-reperfusion injury while restoring cerebral blood flow.The goal of this study is to clarify further part of mechanism of action how QTHY prevents and treats CI/RI.

## 2. Methods

### 2.1 Animal

The Guangxi University of Chinese Medicine’s Laboratory Animal Units offered adult male Sprague-Dawley (SD) rats weighing 250–280g. The experimental scheme, examined by Guangxi University of Chinese Medicine Institutional Animal Welfare and Ethical Committee, matched animal protection, animal welfare ethical principles and relevant rules by National Laboratory Animal Welfare Ethics. The approval number is DW20210511-094.

### 2.2 Preparation of QTHY and TXL

Experimental medicinal materials bought from The Guangxi International Zhuang Medical Hospital were verified as real herbs. They were Tao Ren 10g, Leech 5g, Radix Paeoniae 15g, Salviae Miltiorrhizae 15g, Cyathulae Radix 15g, Chuan Xiong 10g, Di Long 8g, Mao Dongqing 15g, Cortex Moutan 10g, Jiang Banxia 6g, Bile Arisaema 6g. Free-boiled QTHY Decoction is comparable to 5.97g of raw medication per gram, or 1.95g/kg for an adult dose. Tong-xin-luo (TXL) group is comparable to 0.12g/kg for an adult dose. The dosage of oral administration in rats, which originate from combining the clinical dose with conversion coefficient of body surface area from Pharmacological Laboratory Methodology, is gotten.

### 2.3 Determination of therapeutic time window of CI/RI

The rats were divided into blank group (Blank), sham-operated (Sham), cerebral ischemia-reperfusion group (I/R), Qing-tong-hua-yu Decoction group (QTHY) and positive control group (TXL). The TXL and QTHY groups were pretreated 7 days before surgery and given 0.12g/kg Tong-xin-luo and 1.95g/kg Qing-tong-hua-yu Decoction respectively. The Sham group was given with 2ml physiological saline by oral administration for 7 days. After ischemia-reperfusion, each group continued to use original scheme of oral administration. Besides, the corresponding samples were collected at four corresponding time points of 3h, 24h, 3d and 7d after ischemia-reperfusion.

### 2.4 Preparation of rat brain ischemia-reperfusion injury model

To create a rat model of cerebral ischaemia-reperfusion injury (CI/RI) [7, 8], the I/R, TXL, and QTHY groups did CI/RI surgical procedures. The rat was anesthetized by inhaling isoflurane (induction concentration adjusted to 3-4%), maintained a temperature of 37 degrees with a heating pad, and had the right middle cerebral artery blocked for two hours before reperfusion caused transient focal cerebral ischaemia. The Sham group die aame operation, but there was no arterial blockage.

### 2.5 Determination of neurological defect score

The rats’ behavior was examined after reperfusion and graded using the Longa criteria[9]. Inclusion criteria were as follows: A score of 0 means no neurological deficit; A score of 1 means the front paw on the paralyzed side cannot be fully extended; a score of 2 means the walking pattern involves circles directed at the paralyzed side; a score of 3 involves leaning toward the paralyzed side; and A score of 4 means the inability to walk on its own and unconsciousness.

### 2.6 Triphenyltetrazolium chloride (TTC) staining

Rats were executed after the procedure, and the brains were immediately taken out. Coronal sections of 2 mm in thickness were cut from brains. Next, sections were exposed to 2% TTC for 30 min in the dark at 37°C. Then, a scan was done on the stained parts. By using image analysis, the infarct volume was determined and represented as a percentage of the whole brain tissue.

### 2.7 Immunohistochemistry

After dewaxing into water, paraffin sections started antigen repair by citric acid method (G1202, Servicebio). After cooling, endogenous peroxidase was blocked according to the immunohistochemistry kit → blocking → overnight incubation at 4 °C with primary antibody (K001611P, K002209P, Solarbio) → incubation with secondary antibody (G1209, Servicebio) → DAB (G1211, Servicebio) colour development, etc. Finally, the sections were air-dried and sealed, observed under a microscope and photographed, processed by Nikon DS-U3 imaging system, and the average optical density was calculated by using Image-pro plus 6.0 image analysis software.

### 2.8 Western blot analysis

At the designed time point, brain tissue from the anterior 2/3 of the ischemic side of the brain was taken from each group, homogenized on ice with lysis solution (R0010, Solarbio) and centrifuged at 12000 r/min for 5 min at 4°C. After centrifugation, the supernatant was removed and preservated in separate packages. Brain tissue protein concentrations were determined by using a BCA protein concentration assay kit (PC0020, Solarbio) with 4×protein loading buffer (P1016, Solarbio), heated to denature the proteins, separated by 10% SDS polyacrylamide (G2003, Servicebio) gel electrophoresis, and transferred to PVDF membranes by wet transfer. The shaker was closed in a room temperature (5% skimmed milk powder) and washed with EGFR (rat 1:500, Solarbio, K002209P), p44/42MAPK (1:1000, Solarbio, K001611P), GAPDH (1:1000, Bioswamp, PAB45851). After washing, the membrane was placed with the corresponding secondary antibody HRP-labelled goat anti-rabbit IgG (1:1000, Servicebio, GB23303). Finally, the images were developed by gel imaging system (Tanon 5200, Shanghai Tennant), and analysed by using Image J software.

### 2.9 Real-time PCR

Each group of rats had brain tissue samples obtained from the anterior 2/3 of the ischemic side for the mRNA test. Total RNA was extracted from brain tissue with TriQuick Reagent of Solarbio RNeasy reagents according to the manufac-turer’s instruction. A 1 μL RNA sample was taken.Then, the RNA concentration was determined by using a multifunctional enzyme marker and directly determined by A260/280, which represents the purity of the RNA (1.8-2.0). The cDNA was reverse transcribed by using total tissue RNA as template. Primers were designed according to the gene sequence of the gene library and synthesised by Beijing DynaScience Biotechnology Co Ltd (Nanning). The primer sequences are described in Table 1 below. RT-PCR amplification conditions: pre-denaturation 95°C: 30 s; PCR cycles (40 cycles): denaturation: 95°C, 10 s; annealing and extension: 60°C: 30 s. After normalisation to the housekeeping gene GAPDH, the data were quantified by using the 2-ΔΔCt method.

**Table 1.**
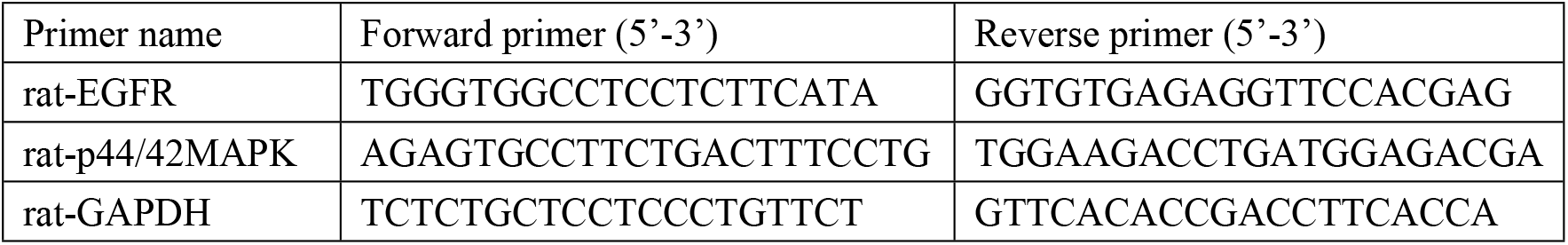
Primer sequences for qPCR

### 2.10 Network construction

#### 2.10.1 Database construction

The “OMIM”[10], “GeneCard”[11], and “DrugBank”[12] databases were used to gather disease-related targets for CI/RI. The TCMSP (https://tcmsp-e.com/index.php) and the Chemistry Database of CAS[13] provide information about QTHY components (Tao Ren, Leech, Radix Paeoniae, Salviae Miltiorrhizae, Cyathulae Radix, Chuan Xiong, Di Long, Mao Dongqing, Cortex Moutan, Jiang Banxia, bile arisaema).

The compounds were screened by oral bioavailability (OB) ≥ 30% and drug-like properties (DL) ≥0.18, which was set according to the chemical composition of TCM; the main active ingredients of QTHY were also supplemented from published papers. The CAS numbers and Pubchem Cid numbers were queried and passed through PubChem[14] to generate 2D protein target structure maps, which were screened for active ingredients by SwissADME[15]. Then, according to Pharmacokinetics, protein targets which have low Gastrointestinal Absorption and low Blood-brain Barrier were excluded. Finally, the Uniprot[16] database was used to convert the target protein names into standard gene names to obtain the genes of the corresponding key targets of the compounds; SwissTargetPrediction[17] was used for target prediction to obtain the target genes corresponding to the QTHY components.

#### 2.10.2 Network construction and analysis

Protein-protein interaction (PPI) networks and drug-compound-acting target-disease networks of overlapping genes were constructed. In order to explore the interactions between the therapeutic targets of Qing-tong-hua-yu Decoction and identify key proteins, the overlapping potential targets were set species to “Multiple proteins” by STRING 11.5 database[18];Meanwhile, Other parameters were default values.The final PPI network was created through the interaction network drawn by using Cytoscape 3.7.2 software, with the node size and color reflecting the magnitude of the degree values and the edge thickness reflecting the size of the aggregate scores.

In order to learn more about the degree of correlation between the main components of QTHY and their corresponding targets of disease, function, and pathway, GO function and KEGG pathway enrichment analyses were carried out on overlapping genes at *P*<0.01 by using online platform of Metascape[19] and the BATMAN-TCM[20]. The top-ranked biological processes and pathways were screened, and the enrichment analysis results were visualized.

### 2.12. Statistical analysis

All data were expressed as means plus or minus standard deviation (X±S) and statistical analysis was performed using SPSS 25.0 software. Comparisons between groups were made using one-way ANOVA and for neurological deficit scores, data were expressed as median and non-parametric statistical analysis was performed by using the Kruskal-Walis test to compare the differences between the two groups. Differences were considered statistically significant at *P*<0.05.

## 3. Results

An eesearch methodology combining pharmacological network analysis and experimental validation was used in this study to uncover the mechanism of action of QTHY in CI/RI. The process, involving the workflow depicted in Figure 1, included five steps: (1) Several typical indicators in the process that QTHY prevent CI/RI; (2) Mining various databases to identify the prescription components and their corresponding disease-related targets; (3) Building a composite target-pathway-disease and function relationship through interaction networks, discovering the most important diseases and functions through network analysis, and identifying the most classical pathways. (4) The potential mechanism of prescriptions in brain protection are evaluated, and the correctness of network analysis was ensured through experiments.

**Fig 1.**
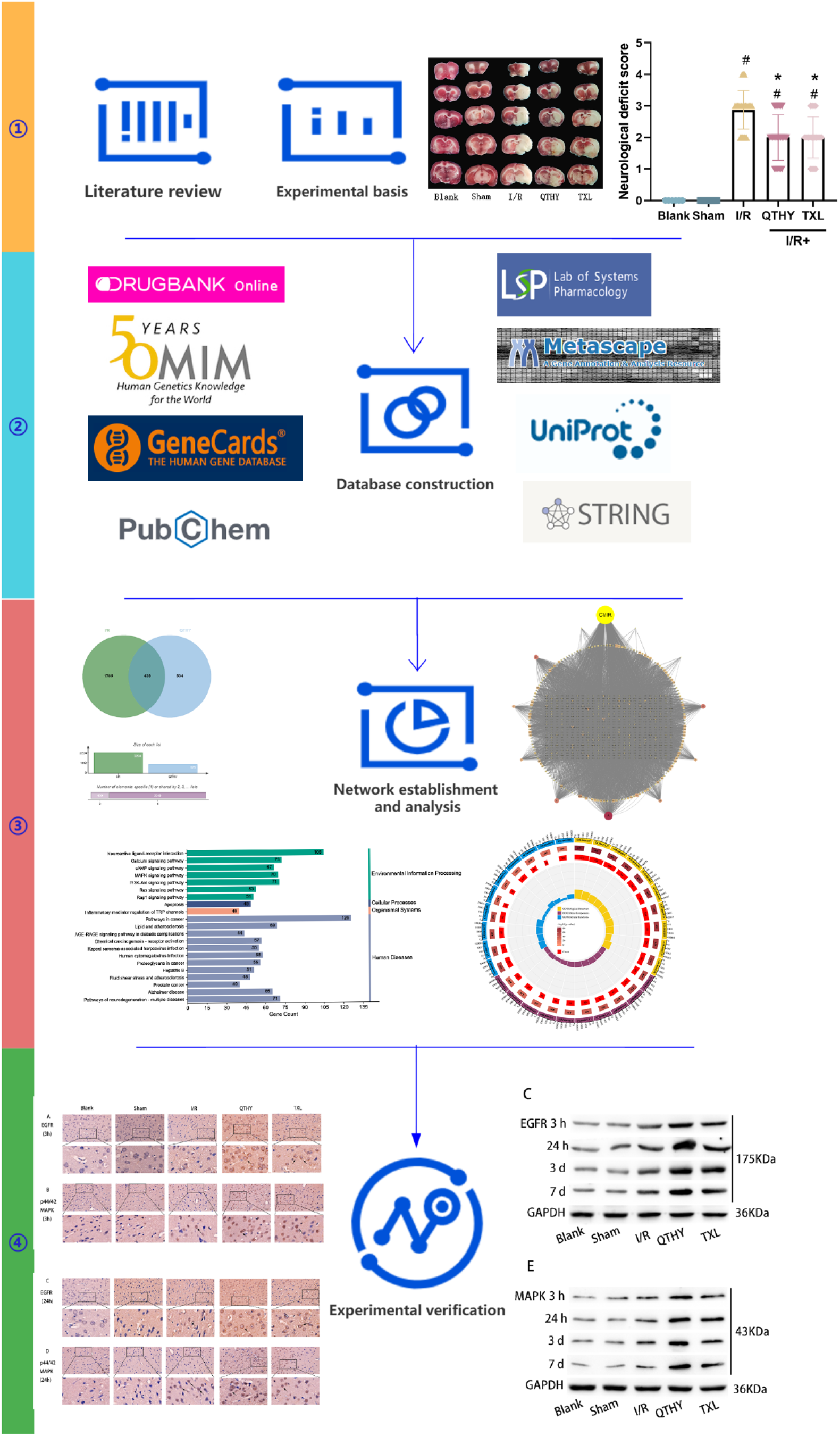
Workflow of QTHY for cerebral protection in the CI / RI model.

### 3.1 QTHY pretreatment can improve neurological deficits

In order to evaluate the protective impact of QTHY on neurological impairments, we performed MCAO with 2h occlusion followed by 3h, 24h, 3d, and 7d reperfusion. To evaluate neurological deficiencies, the Longa scale was applied. Rats that underwent sham surgery moved normally and showed no neurological impairments, as seen in Fig 2. Cerebral ischaemia dramatically changed the neurobehavior of the rats in the model group, as shown by the neurological impairment score. Neurological deficits of rats in the model, QYHY, and TXL groups were significantly worse than those in the sham-operated group at the same time point (*P*<0.05); in contrast, neurological deficits of rats which pretreated with QTHY and the positive control drug TXL were significantly better than those in the model group (*P*<0.05).

**Fig 2.**
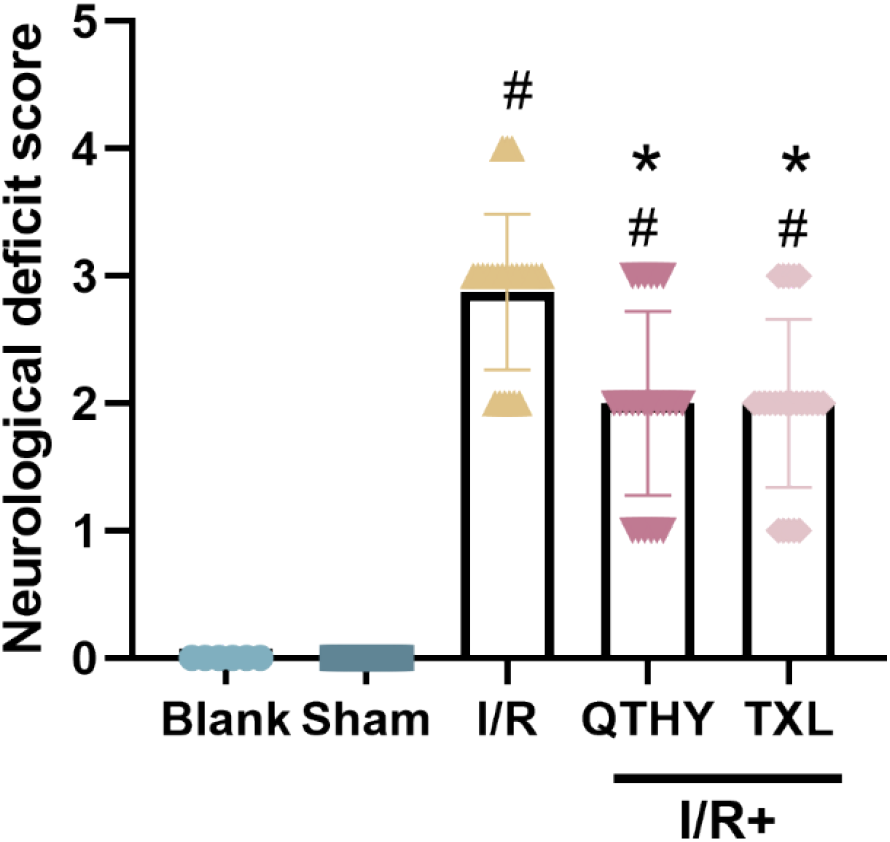
Neurological deficit scores. Neurological deficit score data reported as median, #*P*<0.05 vs sham group; \**P*<0.05 vs I/R group; △*P*<0.05 vs TXL group.

### 3.2 QTHY pretreatment reduces cerebral infarct volume

The protective effect of QTHY pretreatment on infarct volume in MCAO rats was assessed by using TTC staining. TTC-stained brain sections which are representative of each group were selected, as shown in Figure 3A; and corresponding infarct volumes and statistics were got, as shown in Figure 3B. Rats in the sham-operated group had no infarcts. The transient MCAO model group showed a significant increase in infarct volume. Compared with the sham-operated group over the same time period, cerebral infarct volume significantly increased in the model, QTHY and TXL groups (*P*<0.05); Compared with the model group, transient MCAO-induced infarction was significantly inhibited in the QTHY and TXL groups (*P*<0.05).

**Fig 3.**
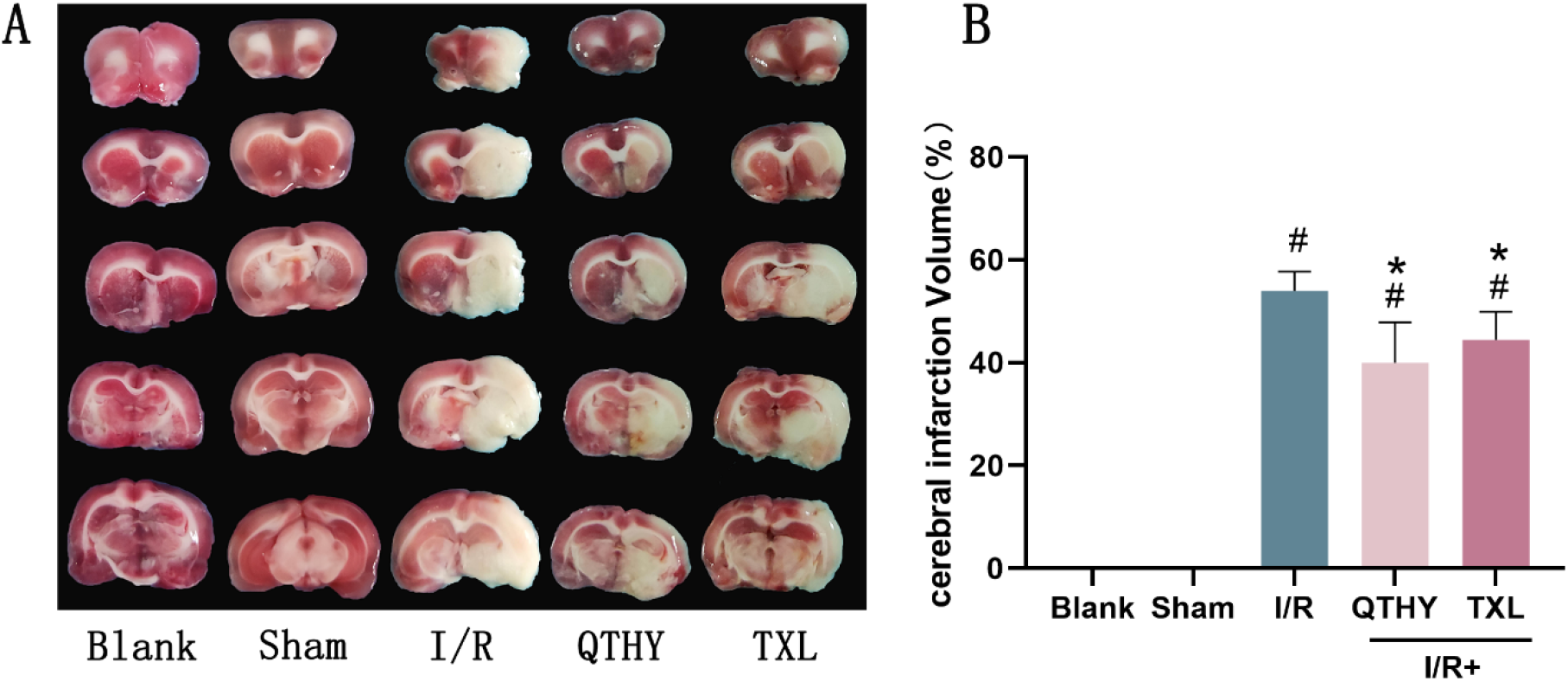
Representative images of TTC-stained brain sections from each group. Infarcted areas are white, while normal areas are red. Quantitative analysis of infarct volumes in each group (n=6). QTHY and TXL treatment significantly reduced infarct volumes, #*P*<0.05 vs sham group; \**P*<0.05 vs I/R group; △*P*<0.05 vs TXL group.

### 3.6 Target disease and functional pathways of QTHY components in CI/RI control

With 439 items in CI/IR associated with QTHY components through Venn intersection (Fig 4), a composite target combined with target disease and functional pathway network was established in order to elucidate the multi-target pharmacological mechanism of QTHY in CI / RI prevention and treatment. Fig 5 visualises the multidimensional interactions between multiple components and multiple targets of Chinese herbal medicine to prevent and treat disease. The results show that beta-sitosterol, palmitoleic acid, palmitic acid, linoleic acid, soy Stigmasterol and other compounds play a key role in the overall network.

**Fig 4.**
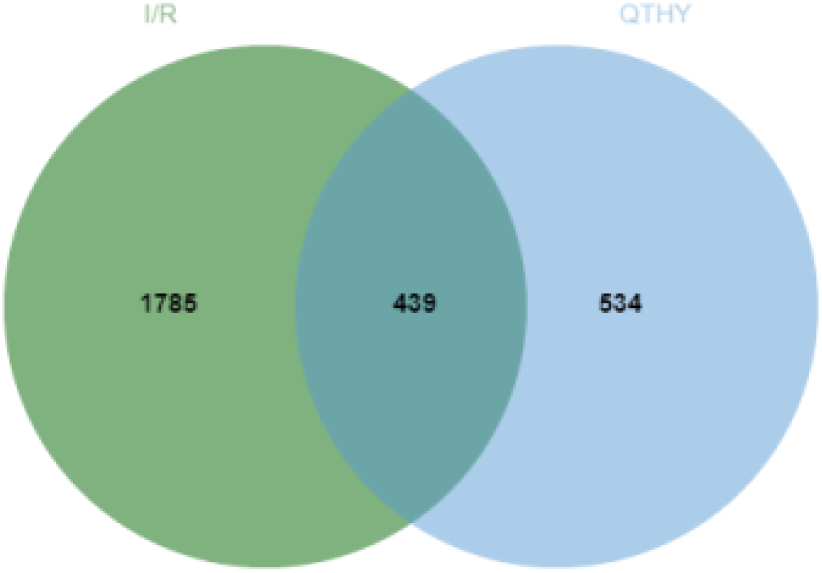
Target intersection of CI/RI and QTHY.

**Fig 5.**
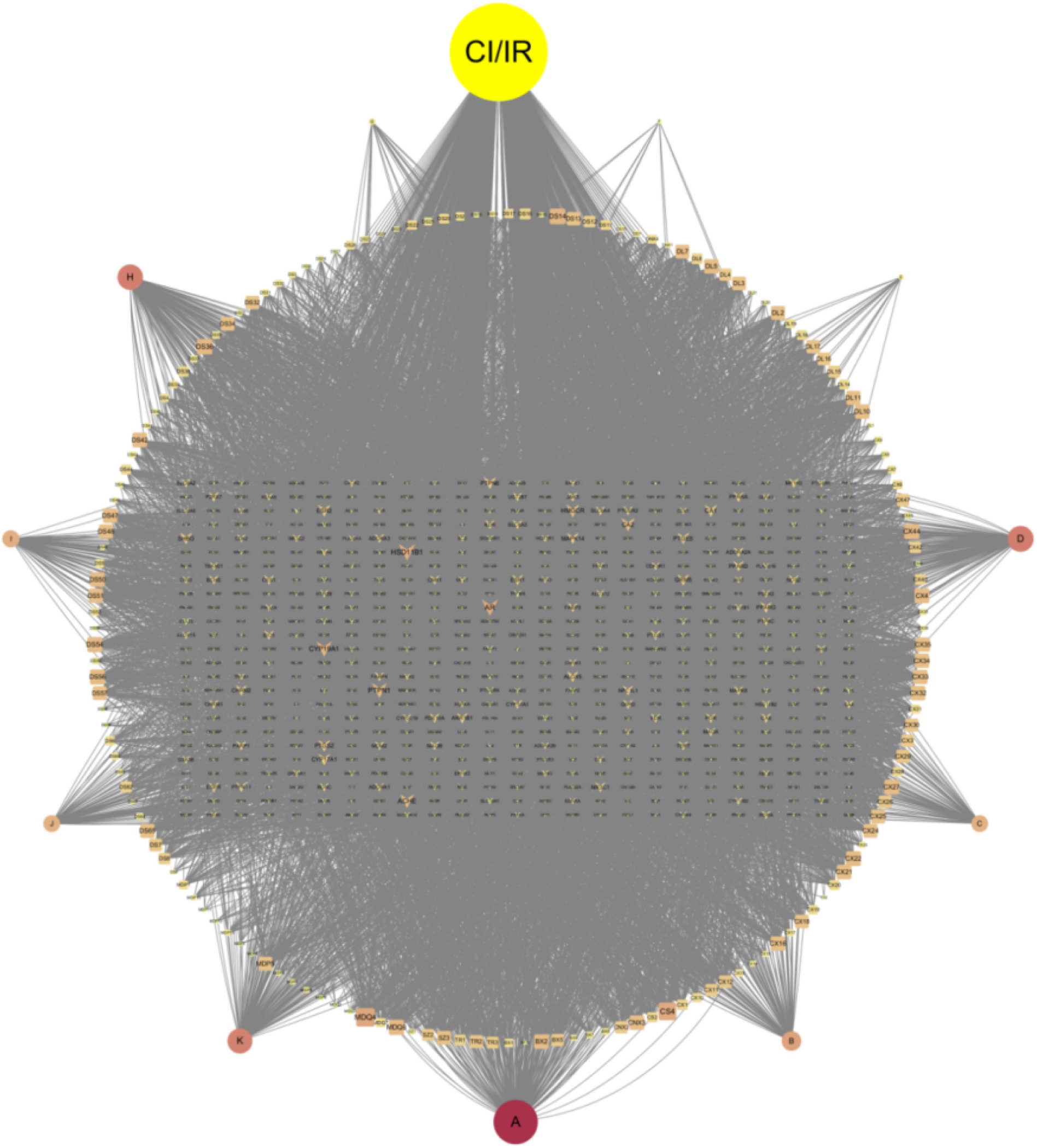
Target network of QTHY component and disease related. The circles represent the QTHY components, with the QTHY component in the middle of the targets associated with 439 CI/RI. Each edge indicates the relationship between compound and target interactions and the core nodes were selected based on network topological features such as node degree values, with the size and shade of colour of the nodes representing the more important role played.

A target-disease and function-pathway network comprising relevant diseases, relevant functions and interconnections was established to elucidate the main biological processes and molecular mechanisms of QTHY action on CI/RI-related targets. GO and KEGG enrichment analyses were performed according to the Metascape and BATMAN-TCM online platform algorithms. The biological process (BP), molecular function (MF) and cellular components (CC), Kegg pathway and disease were ranked according to the corresponding *p* value scores to distinguish the relevance or importance between the 439 targets and multiple diseases, functions and pathways. The target-related pathways most significantly affected by QTHY were ranked in decreasing order of -log (*p* value) score, and the main pathways associated with the target were obtained (Fig 5B): Pathways in cancer, Neuroactive ligand-receptor interaction, Lipid and atherosclerosis, Calcium signaling pathway, cAMP signaling pathway, Human cytomegalovirus infection, MAPK signaling pathway, AGE-RAGE signaling pathway in diabetic complications, Chemical carcinogenesis - receptor activation, PI3K-Akt signaling pathway, Fluid shear stress and atherosclerosis, Kaposi sarcoma-associated herpesvirus infection, Alzheimer disease, Pathways of neurodegeneration - multiple diseases, Proteoglycans in cancer, Hepatitis B, Prostate cancer, Apoptosis, Ras signaling pathway, Pancreatic cancer, EGFR tyrosine kinase inhibitor resistance.

Meanwhile, QTHY could impact a number of functions (Figure 6A): Descending order of -log (*p* value) for BP: cellular response to nitrogen compound, cellular response to organonitrogen compound, regulation of system process, response to inorganic substance, circulatory system process, positive regulation of MAPK cascade, response to hormone, regulation of MAPK cascade, response to molecule of bacterial origin, regulation of ion transport.

**Fig 6.**
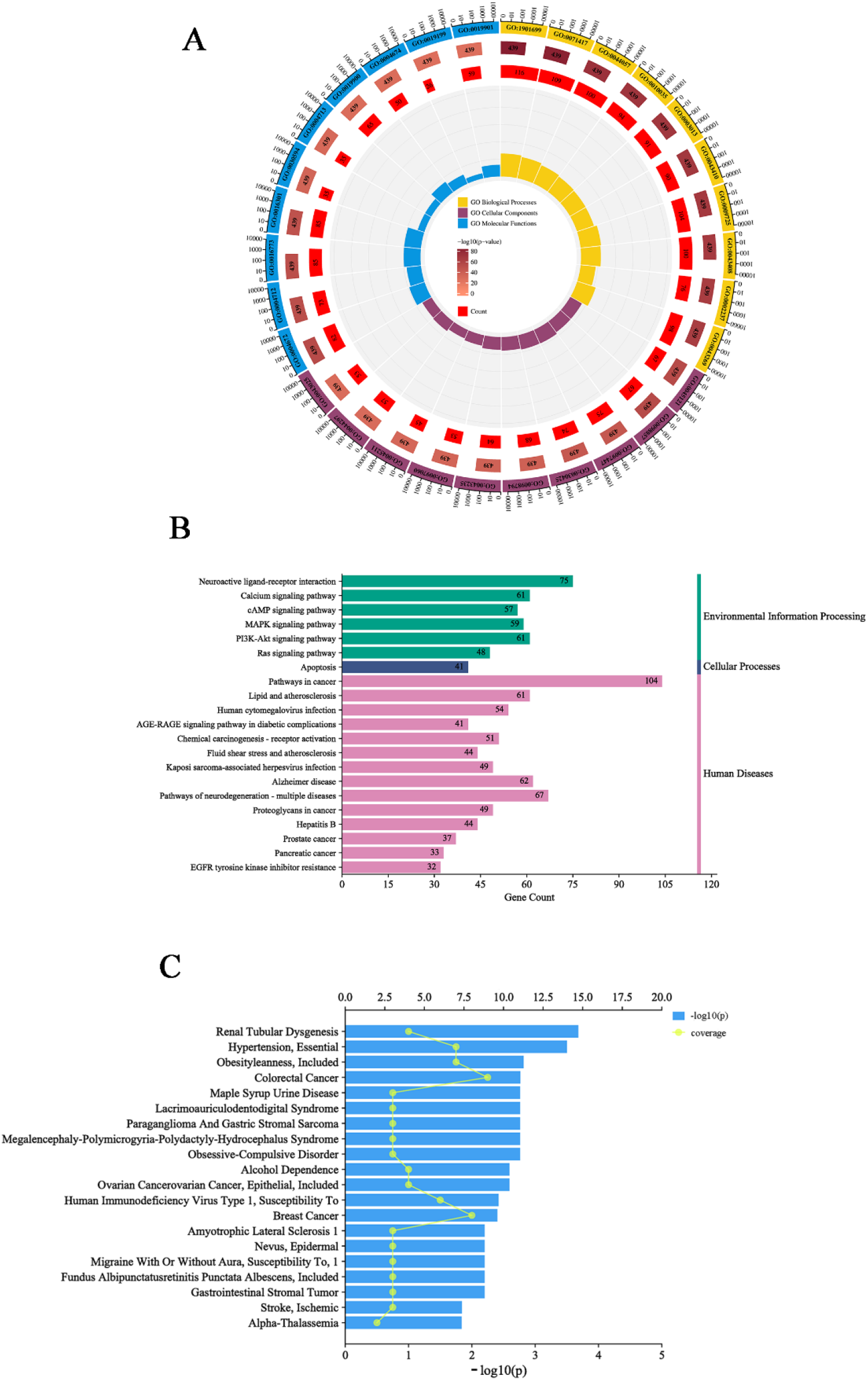
GO and KEGG pathway enrichment analysis of potential targets of QTHY for CI/RI treatment (*P*<0.01). Biological processes, cellular components, molecular functions, functional pathways and diseases affected by QTHY are ranked. The top 10 biological processes, cellular components, molecular functions (A), the top 20 KEGG pathways (B) and diseases affected by QTHY (C) are shown in descending order based on -log (*p* value) scores.

Descending order of -log (*p* value) for MF: protein kinase activity, protein serine/threonine/tyrosine kinase activity, phosphotransferase activity, alcohol group as acceptor, kinase activity, neurotransmitter receptor activity, protein tyrosine kinase activity, kinase binding, protein serine/threonine kinase activity, transmembrane receptor protein kinase activity, protein kinase binding, Descending order of -log (*p* value) for CC: membrane raft, membrane microdomain, dendritic tree, dendrite, postsynapse, receptor complex, synaptic membrane, postsynaptic membrane, cell body, neuronal cell body.The diseases most significantly affected by QTHY are listed in decreasing order of -log (*p* value) score (Fig 6C): Renal Tubular Dysgenesis, Hypertension, Obesityleanness, Colorectal Cancer, Maple Syrup Urine Disease, LADD Syndrome, Paraganglioma And Gastric Stromal Sarcoma, Megalencephaly-Polymicrogyria-Polydactyly-Hydrocephalus Syndrome, Obsessive-Compulsive Disorder, Alcohol Dependence, Ovarian Cancerovarian Cancer, Human Immunodeficiency Virus Type 1, Breast Cancer, Amyotrophic Lateral Sclerosis 1,Nevus, Migraine, Fundus Albipunctatusretinitis Punctata Albescens, Gastrointestinal Stromal Tumor, Ischaemic Stroke and Alpha-Thalassemia.

The primary pathways which are relevant to the targets were obtained based on the results of core and canonical pathway analyses of the 439 targets. QTHY may be able to modify ischaemic stroke either by modifying MAPK cascade responses or through MAPK cascade responses.

### 3.8 Identification of a target network for QTHY regulation of the EGFR/MAPK signalling cascade

In order to determine whether EGFR is a primary target of QTHY effect, a PPI network comprising 439 pertinent targets that were co-regulated by illness and QTHY components was initially created (Fig 7A). In the PPI network, nodes represent proteins, edges represent protein-protein interactions, and node size and colour represent degree values. The higher the degree of a node, the more significant its role in the network. Each edge in the network reflects a protein-protein interaction, whlie more lines indicate a stronger relationship.

**Fig 7.**
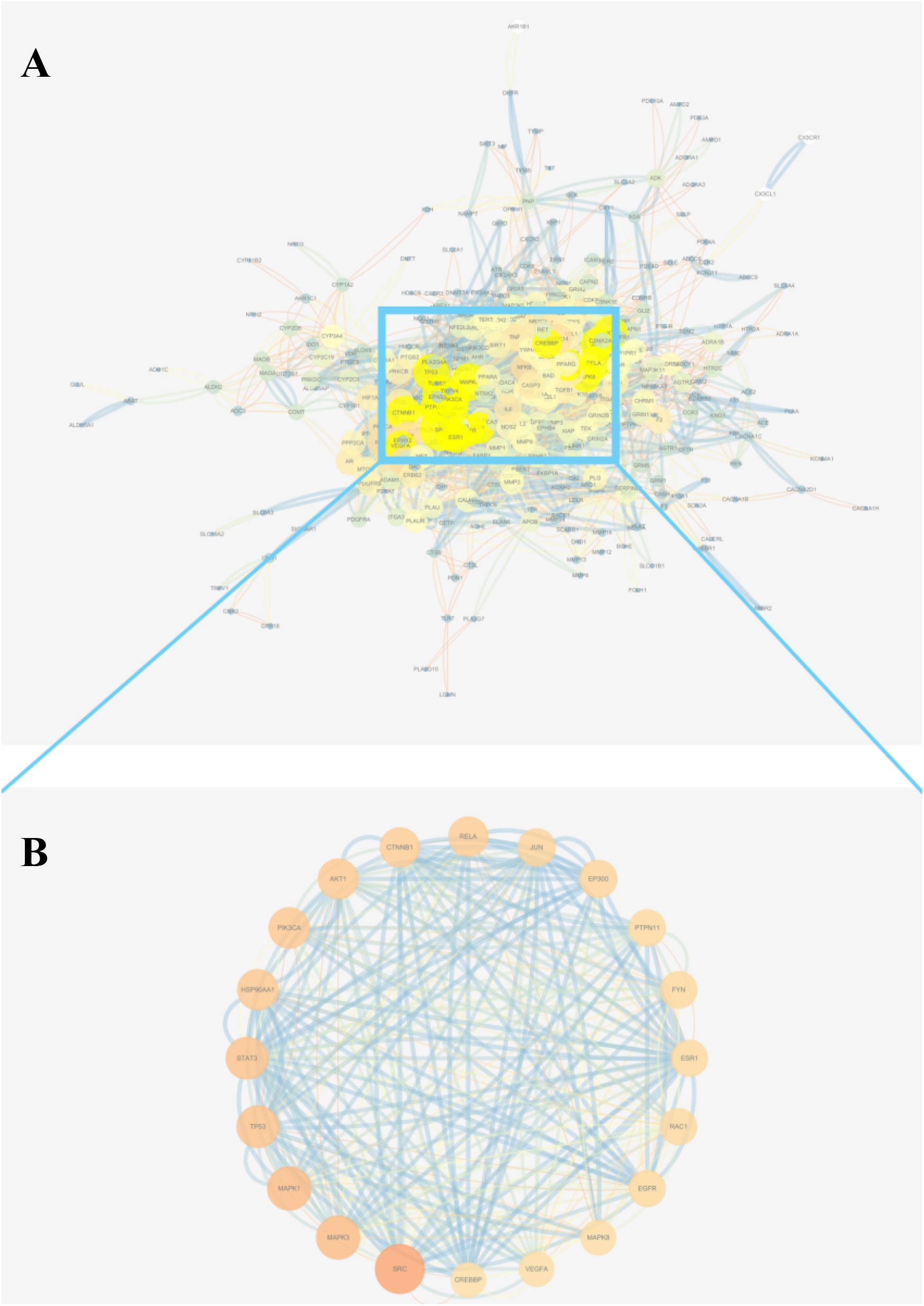
Key targets modulated by QTHY. The top 20 targets are retrieved, and the PPI network’s nodes represent proteins, edges represent protein-protein interactions, and node size and colour represent degree values. The higher the degree of a node, the more significant its role in the network. Each edge in the network reflects a protein-protein interaction, whlie more lines indicate a stronger relationship.

**Fig 8.**
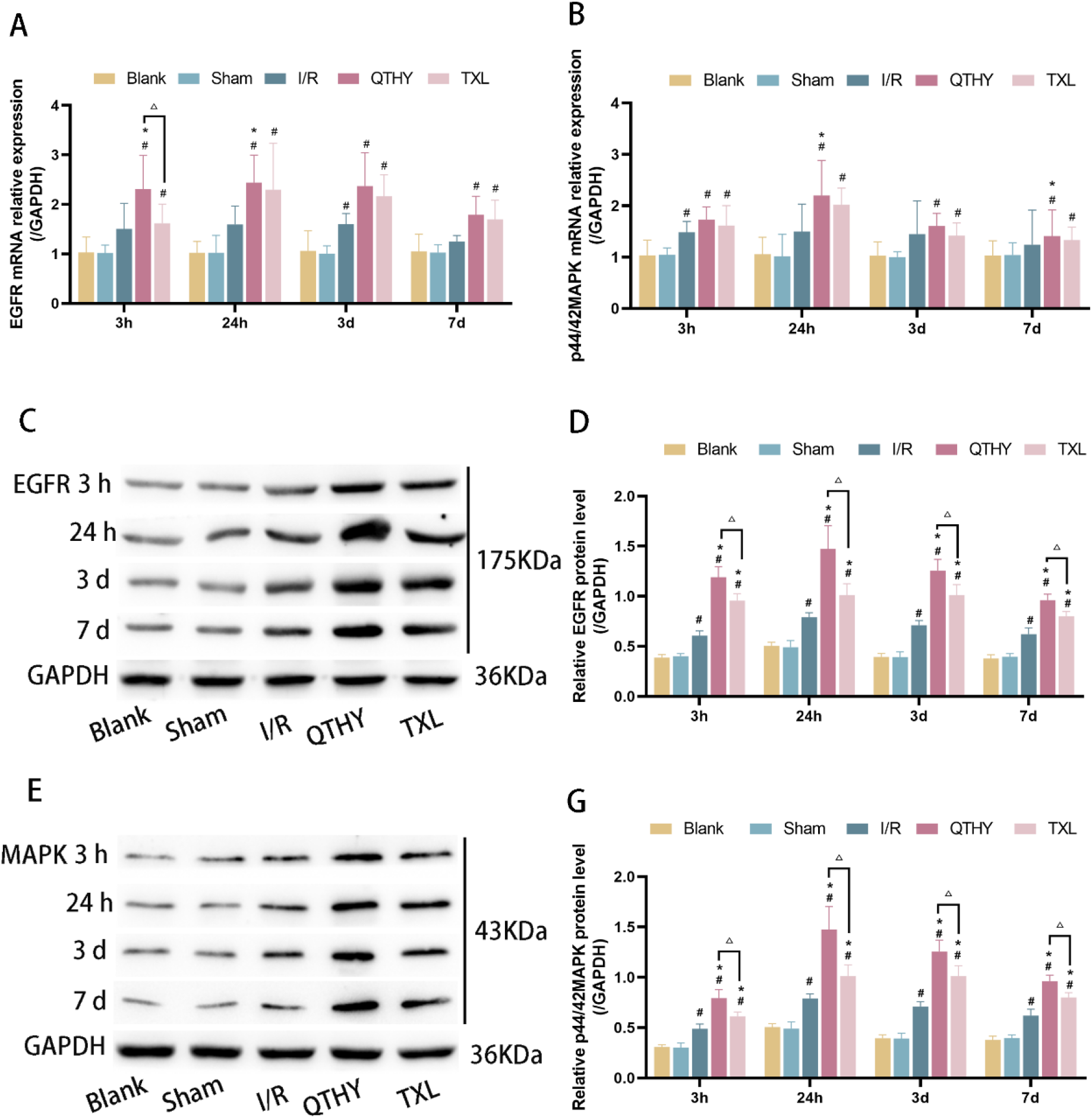
Pretreatment with QTHY modulates the EGFR/MAPK signalling cascade response in the rat brain after CI/RI. EGFR (A), p44/42MAPK (B) mRNA relative expression levels were assessed by RT-qPCR. EGFR, p44/42MAPK in the blank, sham control, I/R, QTHY and TXL groups at each representative protein blot images for the time periods (CE). Statistics of EGFR, p44/42MAPK protein levels in the above four groups (n=6) (DG), normalizing the results to GAPDH. #*P*<0.05 vs sham group; \**P*<0.05 vs I/R group; △*P*<0.05 vs TXL group.

**Fig 9.**
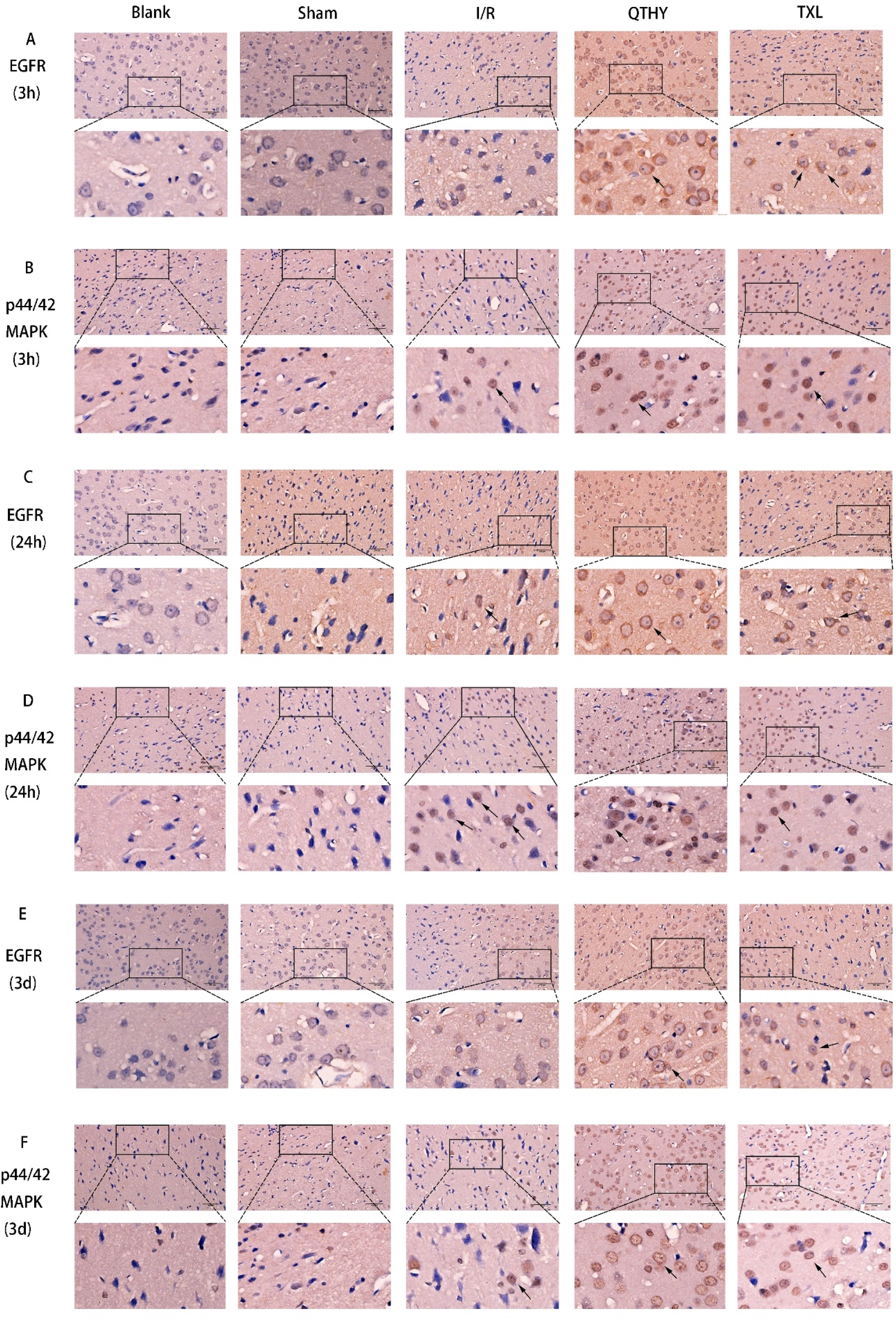
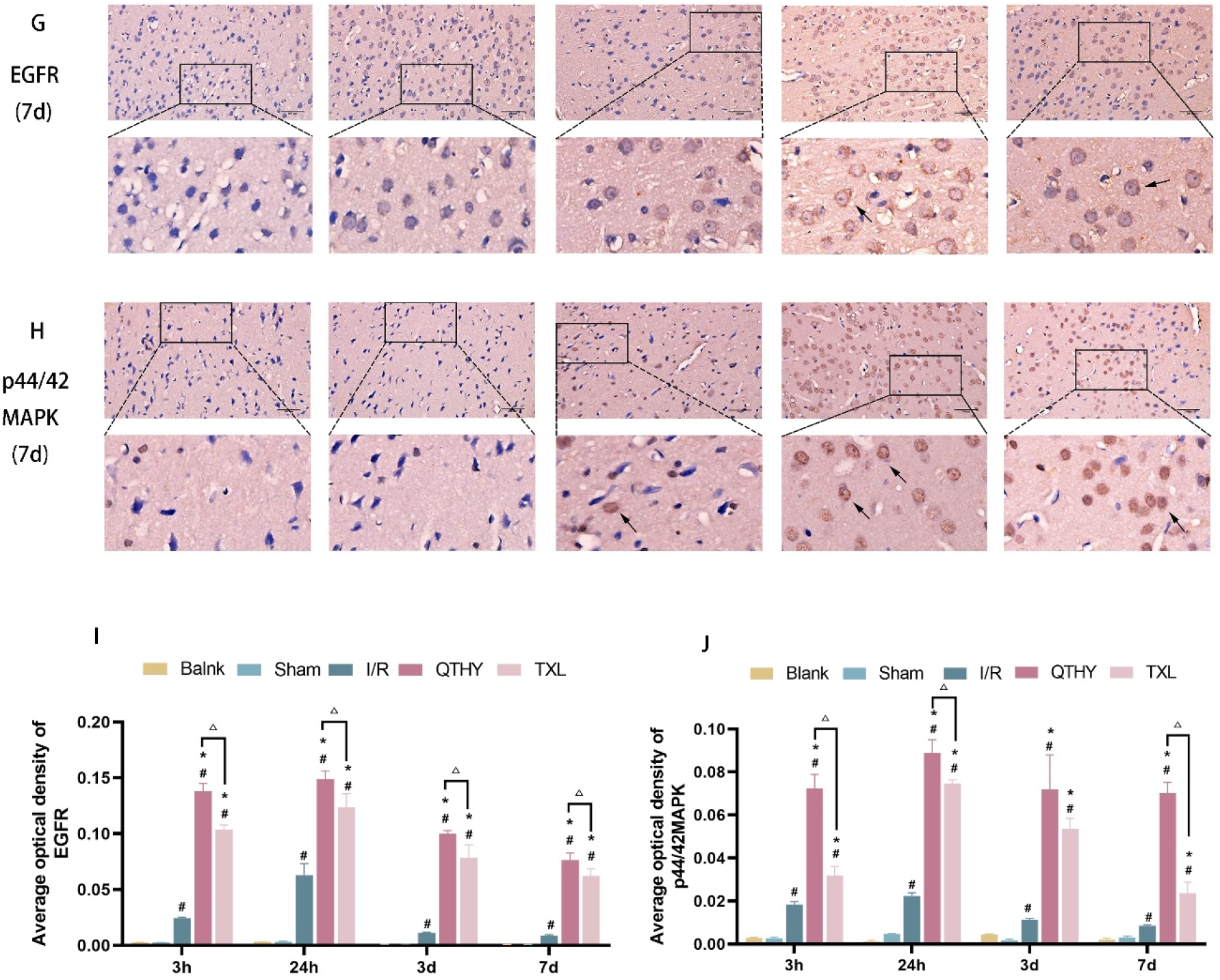
Immunohistochemical staining (×400) of EGFR, p44/42MAPK expression in the SVZ region of rat brain affected by ischemia-reperfusion injury by QTHY (local magnification 40%, black arrows point to coloured cells). (I J) The histogram below shows the mean optical density rate of the affected brain regions in the cerebral cortex under different treatment conditions. #*P*<0.05 vs sham group; \**P*<0.05 vs I/R group; △*P*<0.05 vs TXL group.

To facilitate observation, the top 20 targets in terms of degree value are shown in magnification. As can be seen from Figure 7, the degree values, like SRC、MAPK1、MAPK3、STAT3、TP53、PIK3CA、HSP90AA1、AKT1、RELA、CTNNB1、JUN、EP300、PTPN11、FYN、ESR1、 EGFR、RAC1 、VEGFA 、MAPK8 、CREBBP, were significantly higher than the rest of the targets, which play an important linkage role in the PPI network and may be the most important potential targets of QTHY for CI/RI treatment. The degree value of EGFR was 41 and ranked 16, further demonstrating the important role of EGFR in the protective effect of QTHY against CI/RI. Combined with disease and function analysis, the target of EGFR was considered to be the most critical target in the MAPK cascade response or modulation of the MAPK cascade response.And then the target of EGFR was subsequently selected for further experimental validation.

### 3.8 QTHY is required to regulate the neuroprotective activity of targets in the EGFR/MAPK signalling cascade response

To verify the regulatory effect of QTHY on the above predicted core targets(Fig 6), at the same time point, compared with the sham-operated group, expression levels of p44/42MAPK mRNA were significantly higher in the I/R group 3 hours after reperfusion (*P*<0.05); expression levels of EGFR mRNA were significantly higher 3 days after reperfusion (*P*<0.05); at all time points after reperfusion, expression levels of EGFR and p44/42MAPK mRNA were significantly higher in the QTHY and TXL groups (*P*<0.05); compared with the I/R group, expression levels of EGFR mRNA were significantly higher in the QTHY group 3 hours and 24 hours after reperfusion (*P*<0.05); compared with the I/R group, expression levels of p44/42MAPK mRNA were significantly higher in the QTHY group 24 hours and 7 days after reperfusion (*P*<0.05). At the corresponding time points, the TXL group upregulated the expression of EGFR and p44/42MAPK mRNA to some extent compared with the model and QTHY groups, but none of the differences were statistically significant.

At the same time point, comparison of EGFR and p44/42MAPK proteins in various groups of rats: compared with the sham-operated group, protein expression levels were significantly higher in the I/R, QTHY and TXL groups (*P*<0.05); compared with the I/R group, protein expression levels were significantly higher in the QTHY and TXL groups (*P*<0.05); compared with the TXL group, protein expression levels were significantly higher in the QTHY groups (*P*<0.05).

To better visualize the expression of EGFR and MAPK, we used immunohistochemical staining to show a representative field of view that was selected in each group separately (Fig 7). At the same time point, compared with the sham-operated group, the expression of EGFR and p44/42MAPK in the SVZ was significantly increased in the I/R, QTHY and TXL groups (*P*<0.05). The colour of the cells has deepened and there is significant swelling of the cell morphology. At the same time point, compared with the I/R group, the number of stained cells was significantly higher (*P*<0.05). However, the increase in the number of EGFR and p44/42MAPK stained cell was more notable in the QTHY group (*P*<0.05).

The findings demonstrated that pretreatment with QTHY dramatically elevated expression levels of EGFR and p44/42MAPK, effectively activating the EGFR/MAPK signalling cascade to shield neurons from cerebral ischemia-reperfusion damage.

## 4. Conclusion

Pharmacological or mechanical thrombolysis to restore blood flow is widely recognized as a clinically effective treatment of ischemic stroke[21].However, less than 10% of stroke patients are eligible for conditions for fibrinogen activator therapy, and half of these patients fail to show clinical improvement[22] or even develop secondary neuronal damage caused by CI/RI. Ischemia-reperfusion injury is a common pathological feature of ischaemic stroke, and brain damage caused by ischaemia and reperfusion is a complex pathophysiological process with multiple mechanisms [23].Numerous studies have demonstrated that activated EGFR controls the migration of neural progenitor cells[24, 25] when it is highly expressed in the SVZ area[26, 27]. EGFR’s autophosphorylation initiates the ERK/MAPK route, which leads to DNA synthesis and cell proliferation[28].

Ischemic/hypoxic stimulation activates EGFR in stem/progenitor cells and mediates neurogenesis in experimental ischemic stroke models [29, 30]. Ischemia is a condition in which a region of the body receives inadequate blood flow, which may cause cell death and tissue damage. There is not enough oxygen in the blood, which is known as hypoxia. In experimental ischemic stroke models, it has been demonstrated that ischemia/hypoxic stimulation activates the EGFR (epidermal growth factor receptor) in stem/progenitor cells, and that this activation mediates neurogenesis (the production of new neurons). This shows that ischemia/hypoxia may contribute to the repair of damaged tissue and the growth of new neurons after stroke[31].MAPK contributes to the maintenance of neural stem cells[32], and can respond to cerebral ischemia by phosphorylation reactions of threonine and tyrosine residues[33]. MAPK phosphorylation promotes neuronal survival in the dentate gyrus area[34].Targeting the EGFR/MAPK signalling cascade response is an important therapeutic goals, which is used to stimulate neurogenesis and brain regeneration for treating post-stroke injuries. The EGFR/MAPK signaling cascade controls cell migration, differentiation, and proliferation which are necessary for neurogenesis and brain regeneration[35]. Thus, the EGFR/MAPK signalling cascade response is an important therapeutic target in the activation of neurogenesis and brain restoration in the treatment of post-stroke dysfunction[35].

The study of the relationships among medications, targets, and illnesses in biological networks is the focus of network pharmacology. Understanding the potentially intricate links between herbal components and ailments might help focus an extensive and methodical investigation of network pharmacology, which is congruent with a holistic viewpoint[36-38]. In recent years, network pharmacology analysis has revealed synergistic effects of multi-component and multi-target drugs and potential pharmacological mechanisms in herbal formulations, and it has also shown that herbal prescriptions have synergistic effects in achieving a cure or reducing toxicity. We use Qing-tong-hua-yu Decoction (QTHY), a clinically effective Chinese herbal formula (CMF), to treat ischaemic stroke because it has the ability to clear cerebral veins and protect the brain from toxicity. It also has a positive impact on lowering the TCM symptom score and neurological deficits in ischaemic stroke. In the current study, similar analysis of QTHY which contains various herbal components revealed a network of complicated target diseases and potential functional pathways, which is complex but still synergistic.We discovered that the identified components of QTHY may affect 439 CI/RI-related targets. A conclusion we obtained in the current study showed that QTHY pretreatment can be neuroprotective by lowering the extent of acute cerebral infarction, supported by the results of an in vivo transient ischemia model in rats. The results of the network pharmacology study showed that QTHY may be EGFR promoter of inducing neurogenesis in post-ischaemic brain damage, and further verified that the EGFR/MAPK signalling cascade plays a key role in ischaemic stroke.

To confirm the reliability of the network pharmacological analysis, we also performed RT-qPCR to detect the mRNA expression of EGFR and p44/42MAPK, protein expression levels of EGFR and p44/42MAPK by protein immunoblotting, and positive expression rate of EGFR and p44/42MAPK by immunohistochemistry. The results of the study showed that the mRNA expression levels of EGFR and p44/42MAPK in brain tissue were elevated in rats pretreated with QTHY compared with the sham-operated group, with a significant increase in protein expression and a significantly higher positive expression rate. As expected, EGFR was actively expressed in areas experiencing neural injury, including the subventricular zone (SVZ) of the lateral ventricles, the granular layer of the dentate gyrus (DG) of the hippocampus of memory, and the cerebellar granular layer[39, 40]. It has been shown that when EGFR is knocked out, mice exhibit defects in cortical neurogenesis and retinal histogenesis[41]. Reduced cell proliferation in the SVZ was observed in another EGFR ligand, TGF-a knockout mice, further suggesting a role for EGFR in cell proliferation[42]. After focal ischemic injury, EGFR expression was significantly increased in the SVZ[3], with cell proliferation enhancing. In the current study, we found that EGFR expression in the SVZ after injury was enhanced, suggesting that the EGFR-mediated signaling cascade is mainly involved in promoting neurogenesis[43]. And EGFR may stimulate G0 quiescent cells to enter S phase[44]. EGFR transfers extracellular signals into cells and induces cell proliferation[45]. In an animal model of experimental ischemic stroke, activated EGFR induces neural precursor cells proliferate in vivo[4, 46] and reduces brain damage[47, 48]. Thus, EGFR is also a signal for cell survival[49, 50].In a word, these studies point that EGF and its receptors are important in potentially influencing cell proliferation both during normal development and after brain injury. Meanwhile, other studies have shown that EGFR-mediated signaling contributes to the transformation of cells into a neuroglial phenotype[51]. In addition, under certain circumstances, EGFR can mediate an EGFR/MAPK signalling cascade response to promote neural stem cell (NSC) proliferation[40]. Activated but not inhibited MAPK signalling cascade has a neuroprotective effect[1, 52], suggesting a role for EGFR expression in MAPK signalling cascade activation to promote neuronal survival.

In conclusion, the brain microenvironment after ischemic injury is very complex. The study shows that QTHY is further confirmed that it has complex pharmacological effects, which may affect interaction of complex environmental factors in the brain and brain repair in transient ischaemic brain areas, resulting in improved neurological function. When cerebral ischemia-reperfusion injury (CI/RI) occurs, QTHY Plays a partial regulatory role through the EGFR/MAPK signalling cascade response, producing a marked effect in neuroprotection and brain repair and improving brain infarct volume in rats with cerebral ischemia-reperfusion damage.

## Author Contributions

**Conceptualization:** Jiajing Hu

**Data Curation:** Long Zuo, Wenyu Qu

**Investigation:** Jiajing Hu, Jie Bao, Wenyan Zhang

**Supervision:** Meizhen Zhu, Tian Li

**Methodology:** Hongdun He, Yunyang Zhang

**Project Administration:** Meizhen Zhu

**Visualization:** Jiajing Hu; Long Zuo, Wenyu Qu

**Writing – Review & Editing:** Jiajing Hu, Long Zuo, Wenyu Qu

## Supporting information

**S1 Raw images**.

(RAR)

**S1 Data**.

(RAR)

## References

1. Teertam SK, Prakash Babu PJSr. Differential role of SIRT1/MAPK pathway during cerebral ischemia in rats and humans. 2021;11(1):1–14.

2. Zhang M, Gong J-X, Wang J-L, Jiang M-Y, Li L, Hu Y-Y, et al. p38 MAPK participates in the mediation of GLT-1 up-regulation during the induction of brain ischemic tolerance by cerebral ischemic preconditioning. 2017;54(1):58–71.

3. Ninomiya M, Yamashita T, Araki N, Okano H, Sawamoto KJNl. Enhanced neurogenesis in the ischemic striatum following EGF-induced expansion of transit-amplifying cells in the subventricular zone. 2006;403(1-2):63–7.

4. Türeyen K, Vemuganti R, Bowen KK, Sailor KA, Dempsey RJJN. EGF and FGF-2 infusion increases post-ischemic neural progenitor cell proliferation in the adult rat brain. 2005;57(6):1254–63.

5. Uzun G, Subhani D, Amor SJJov, neurology i. Trophic factors and stem cells for promoting recovery in stroke. 2010;3(1):3.

6. Sütterlin P, Williams EJ, Chambers D, Saraf K, von Schack D, Reisenberg M, et al. The molecular basis of the cooperation between EGF, FGF and eCB receptors in the regulation of neural stem cell function. 2013;52:20–30.

7. Dong B, Yang Y, Zhang Z, Xie K, Su L, Yu YJBa. Hemopexin alleviates cognitive dysfunction after focal cerebral ischemia-reperfusion injury in rats. 2019;19(1):1–9.

8. Ye Z, Guo Q, Xia P, Wang N, Wang E, Yuan YJBr. Sevoflurane postconditioning involves an up-regulation of HIF-1α and HO-1 expression via PI3K/Akt pathway in a rat model of focal cerebral ischemia. 2012;1463:63–74.

9. Longa EZ, Weinstein PR, Carlson S, Cummins RJs. Reversible middle cerebral artery occlusion without craniectomy in rats. 1989;20(1):84–91.

10. Rasmussen SA, Hamosh AJGiM. What’s in a name? Issues to consider when naming Mendelian disorders. 2020;22(10):1573–5. Database: OMIM [Internet]. Available from: https://omim.org/

11. Safran M, Rosen N, Twik M, BarShir R, Stein TI, Dahary D, et al. The genecards suite. Practical guide to life science databases: Springer; 2021. p. 27–56. Database: GeneCards [Internet]. Available from: https://www.genecards.org/

12. Wishart DS, Feunang YD, Guo AC, Lo EJ, Marcu A, Grant JR, et al. DrugBank 5.0: a major update to the DrugBank database for 2018. 2018;46(D1):D1074–D82. Database: DrugBank [Internet]. Available from: https://go.drugbank.com/

13. Shanghai Institute of Organic Chemistry of CAS. Chemistry Database of CAS[DB/OL]. Database: Chemistry Database of CAS [Internet]. Available from: http://www.organchem.csdb.cn.

14. Kim S, Chen J, Cheng T, Gindulyte A, He J, He S, et al. PubChem in 2021: new data content and improved web interfaces. 2021;49(D1):D1388–D95. Database: PubChem [Internet]. Available from: https://pubchem.ncbi.nlm.nih.gov/

15. Daina A, Michielin O, Zoete VJSr. SwissADME: a free web tool to evaluate pharmacokinetics, drug-likeness and medicinal chemistry friendliness of small molecules. 2017;7(1):1–13. Database: SwissADME [Internet]. Available from: http://www.swissadme.ch/

16. Breuza L, Arighi CN, Argoud-Puy G, Casals-Casas C, Estreicher A, Famiglietti ML, et al. A coordinated approach by public domain bioinformatics resources to aid the fight against Alzheimer’s disease through expert curation of key protein targets. 2020;77(1):257–73. Database: Uniprot [Internet]. Available from: https://www.uniprot.org/

17. Daina A, Michielin O, Zoete VJNar. SwissTargetPrediction: updated data and new features for efficient prediction of protein targets of small molecules. 2019;47(W1):W357-W64. Database: SwissTargetPrediction [Internet]. Available from: http://www.swisstargetprediction.ch/

18. Szklarczyk D, Gable AL, Nastou KC, Lyon D, Kirsch R, Pyysalo S, et al. The STRING database in 2021: customizable protein–protein networks, and functional characterization of user-uploaded gene/measurement sets. 2021;49(D1):D605–D12. Database: STRING [Internet]. Available from: https://cn.string-db.org/

19. Zhou Y, Zhou B, Pache L, Chang M, Khodabakhshi AH, Tanaseichuk O, et al. Metascape provides a biologist-oriented resource for the analysis of systems-level datasets. 2019;10(1):1–10. Database: Metascape [Internet]. Available from: https://metascape.org/

20. Liu Z, Guo F, Wang Y, Li C, Zhang X, Li H, et al. BATMAN-TCM: a bioinformatics analysis tool for molecular mechANism of traditional Chinese medicine. 2016;6(1):1–11. Database: BATMAN-TCM [Internet]. Available from: http://bionet.ncpsb.org.cn/

21. Gao J, Long L, Xu F, Feng L, Liu Y, Shi J, et al. Icariside II, a phosphodiesterase 5 inhibitor, attenuates cerebral ischaemia/reperfusion injury by inhibiting glycogen synthase kinase-3β-mediated activation of autophagy. 2020;177(6):1434–52.

22. Tao T, Liu M, Chen M, Luo Y, Wang C, Xu T, et al. Natural medicine in neuroprotection for ischemic stroke: challenges and prospective. 2020;216:107695.

23. Lin L, Wang X, Yu ZJB, access po. Ischemia-reperfusion injury in the brain: mechanisms and potential therapeutic strategies. 2016;5(4).

24. Caric D, Raphael H, Viti J, Feathers A, Wancio D, Lillien L. EGFRs mediate chemotactic migration in the developing telencephalon. 2001.

25. Abe Y, Nawa H, Namba HJNr. Activation of epidermal growth factor receptor ErbB1 attenuates inhibitory synaptic development in mouse dentate gyrus. 2009;63(2):138–48.

26. Novak U, Walker F, Kaye AJJocn. Expression of EGFR-family proteins in the brain: role in development, health and disease. 2001;8(2):106–11.

27. Weickert CS, Webster MJ, Colvin SM, Herman MM, Hyde TM, Weinberger DR, et al. Localization of epidermal growth factor receptors and putative neuroblasts in human subependymal zone. 2000;423(3):359–72.

28. Oda K, Matsuoka Y, Funahashi A, Kitano HJMsb. A comprehensive pathway map of epidermal growth factor receptor signaling. 2005;1(1):2005.0010.

29. Alagappan D, Lazzarino DA, Felling RJ, Balan M, Kotenko SV, Levison SWJAn. Brain injury expands the numbers of neural stem cells and progenitors in the SVZ by enhancing their responsiveness to EGF. 2009;1(2):AN20090002.

30. Aguirre A, Rubio ME, Gallo VJN. Notch and EGFR pathway interaction regulates neural stem cell number and self-renewal. 2010;467(7313):323–7.

31. Gu D-m, Lu P-H, Zhang K, Wang X, Sun M, Chen G-Q, et al. EGFR mediates astragaloside IV-induced Nrf2 activation to protect cortical neurons against in vitro ischemia/reperfusion damages. 2015;457(3):391–7.

32. Campos LS, Leone DP, Relvas JB, Brakebusch C, Fässler R, Suter U, et al. β1 integrins activate a MAPK signalling pathway in neural stem cells that contributes to their maintenance. 2004.

33. Zhan L, Yan H, Zhou H, Sun W, Hou Q, Xu EJMn. Hypoxic preconditioning attenuates neuronal cell death by preventing MEK/ERK signaling pathway activation after transient global cerebral ischemia in adult rats. 2013;48(1):109–19.

34. Hu BR, Wieloch TJJon. Tyrosine phosphorylation and activation of mitogen-activated protein kinase in the rat brain following transient cerebral ischemia. 1994;62(4):1357–67.

35. Chen X, Wu H, Chen H, Wang Q, Xie X-j, Shen Jjmn. Astragaloside VI promotes neural stem cell proliferation and enhances neurological function recovery in transient cerebral ischemic injury via activating EGFR/MAPK signaling cascades. 2019;56(4):3053–67.

36. Yuan H, Ma Q, Cui H, Liu G, Zhao X, Li W, et al. How can synergism of traditional medicines benefit from network pharmacology? 2017;22(7):1135.

37. Liu K, Tao X, Su J, Li F, Mu F, Zhao S, et al. Network pharmacology and molecular docking reveal the effective substances and active mechanisms of Dalbergia Odoriferain protecting against ischemic stroke. 2021;16(9):e0255736.

38. Yang H-Y, Liu M-L, Luo P, Yao X-S, Zhou HJP. Network pharmacology provides a systematic approach to understanding the treatment of ischemic heart diseases with traditional Chinese medicine. 2022:154268.

39. Cameron HA, Hazel TG, McKay RDJJon. Regulation of neurogenesis by growth factors and neurotransmitters. 1998;36(2):287–306.

40. Tucker MS, Khan I, Fuchs-Young R, Price S, Steininger TL, Greene G, et al. Localization of immunoreactive epidermal growth factor receptor in neonatal and adult rat hippocampus. 1993;631(1):65–71.

41. Close JL, Liu J, Gumuscu B, Reh TAJG. Epidermal growth factor receptor expression regulates proliferation in the postnatal rat retina. 2006;54(2):94–104.

42. Wong RJC, Cmls MLS. Transgenic and knock-out mice for deciphering the roles of EGFR ligands. 2003;60(1):113–8.

43. Morita M, Kozuka N, Itofusa R, Yukawa M, Kudo YJJon. Autocrine activation of EGF receptor promotes oscillation of glutamate-induced calcium increase in astrocytes cultured in rat cerebral cortex. 2005;95(3):871–9.

44. Ahsan A, Hiniker SM, Davis MA, Lawrence TS, Nyati MKJCr. Role of cell cycle in epidermal growth factor receptor inhibitor-mediated radiosensitization. 2009;69(12):5108–14.

45. Lui V-Y, Grandis JRJAr. EGFR-mediated cell cycle regulation. 2002;22(1A):1–11.

46. Teramoto T, Qiu J, Plumier J-C, Moskowitz MAJTJoci. EGF amplifies the replacement of parvalbumin-expressing striatal interneurons after ischemia. 2003;111(8):1125–32.

47. Justicia C, Planas AMJJoCBF, Metabolism. Transforming growth factor-α acting at the epidermal growth factor receptor reduces infarct volume after permanent middle cerebral artery occlusion in rats. 1999;19(2):128–32.

48. Sun D, Bullock MR, Altememi N, Zhou Z, Hagood S, Rolfe A, et al. The effect of epidermal growth factor in the injured brain after trauma in rats. 2010;27(5):923–38.

49. De la Rosa EJ, de Pablo FJTin. Cell death in early neural development: beyond the neurotrophic theory. 2000;23(10):454–8.

50. Honarpour N, Tabuchi K, Stark J, Hammer RE, Südhof T, Parada L, et al. Embryonic neuronal death due to neurotrophin and neurotransmitter deprivation occurs independent of Apaf-1. 2001;106(2):263–74.

51. Lillien L, Wancio DJM, Neuroscience C. Changes in epidermal growth factor receptor expression and competence to generate glia regulate timing and choice of differentiation in the retina. 1998;10(5-6):296–308.

52. Yang Y, Zhang S, Fan C, Yi W, Jiang S, Di S, et al. Protective role of silent information regulator 1 against hepatic ischemia: effects on oxidative stress injury, inflammatory response, and MAPKs. 2016;20(5):519–31.

